# Pygmaeomycetaceae, a new root associated family in Mucoromycotina from the pygmy pine plains

**DOI:** 10.1101/2020.07.03.187096

**Authors:** Emily Walsh, Jing Luo, Swapneel Khiste, Adam Scalera, Sana Sajjad, Ning Zhang

## Abstract

A new genus, *Pygmaeomyces*, and two new species are described based on phylogenetic analyses, phenotypic and ecological characters. The species delimitation was based on concordance of gene genealogies. The *Pygmaeomyces* cultures were isolated from the roots of mountain laurel (*Kalmia latifolia*) and pitch pine (*Pinus rigida*) from the acidic and oligotrophic New Jersey Pygmy Pine Plains; however, they likely have a broader distribution because their internal transcribed spacer (ITS) sequences have high similarity with a number of environmental sequences from multiple independent studies. Based on the phylogeny and phenotypical characters, a new family Pygmaeomycetaceae is proposed to accommodate this new lineage in Mucoromycotina. Pygmaeomycetaceae corresponds to Clade GS23, which was identified based on a sequence-only soil fungal survey and was believed to be a distinct new class. Compared to the culture-based methods, we observed that sequence-only analyses tend to over-estimate the taxonomic level. Results from this work will facilitate ecological and evolutionary studies on root-associated fungi.

## INTRODUCTION

The New Jersey Pine Plains are a drought and fire-prone ecosystem with acidic and oligotrophic soils embedded within the New Jersey Pine Barrens (Tedrow 1952; Forman 1998; Ledig et al. 2013). They are comprised of four areas, the East Plains, the West Plains, the Little Plains and the Spring Hill Plains, which together form the largest acreage, 12,400 acres (4,950 km^2^), of dwarf pitch pine in the world. While the pine barrens are dominated by pitch pines (*Pinus rigida*) with heights of 4.6-12 m, the vegetation in these plains areas are primarily composed of pitch pines of low stature (<3.3 m), scrub oak (*Quercus ilicifolia*), and an increased occurrence of low shrub species such as mountain laurel (*Kalmia latifolia*), pyxie moss (*Pyxidanthera barbulata*), bearberry, and other members of the heath family (Ericaceae) (Good et al. 1979; Ledig et al. 2013; McCormick 1979). Noticeably absent from these plains are common pine barrens tree species such as black oak (*Quercus velutina*), white oak (*Quercus alba*), scarlet oak (*Quercus coccinea*), chestnut oak (*Quercus prinus*), and shortleaf pine (*Pinus echinata*) (Forman 1998; Good et al. 1979). Published hypotheses of why the pygmy pines have dwarf stature and crooked form vary from soil chemistry, soil physical property, to water table levels and fire frequency (Harshberger 1916; Lutz 1934; Tedrow 1952; Good et al. 1979). The microbial hypothesis was also proposed by Harshberger (1916) over a century ago, who observed that plants grew differently in the heat sterilized Pine Plains soil compared to the untreated soil. However, no further investigation had been done in this area (Harshberger 1916; Andresen 1959). Our results on plant-associated fungi reported in this paper may shed light on the role of these microbial symbionts in the evolution of pygmy pines.

The diversity of fungi and their functions in the acidic, oligotrophic pine barrens ecosystem still remains largely unknown (Tuininga et al. 2004; Forman 1998). Our previous work in the New Jersey Pine Barrens uncovered a number of novel fungal lineages from the roots of Poaceae grasses and *Pinus* trees (Luo et al. 2017). To date, three new genera and over ten new species in Ascomycota have been described, some of which showed a positive or negative effect to plant health while others had no significant effect (Luo and Zhang 2013; Luo et al. 2014b; Walsh et al. 2014; Walsh et al. 2015).

In this study, we uncovered a new genus *Pygmaeomyces* and two new species, *P. thomasii* and *P. pinuum* in Mucoromycotina, isolated from apparently healthy mountain laurel and pygmy pitch pine roots from the acidic Pygmy Pine Plains. They can be distinguished from *Umbelopsis* species and other related fungi based on phylogenetic analyses, ecology and morphological characters. Based on phylogeny and the rate of rDNA sequence divergence, we propose a new family Pygmaeomycetaceae to accommodate this new lineage. We also conducted plant-fungal interaction experiments and enzymatic tests to further understand their roles in the ecosystem. The DNA barcode sequences of the new taxa match with a number of environmental sequences from acidic soils in a worldwide geographical distribution.

## MATERIALS AND METHODS

### Fungal isolation

Roots of mountain laurel (*Kalmia latifolia*) and pygmy pitch pine (*Pinus rigida*) were collected from a dwarf forest (39.80725, −74.404350 elev. 194ft.) in the Pygmy Pine Plains in New Jersey, USA in July 2016. The soil of the sampling location was acidic (pH 4.1), with low phosphorous (3 ppm), low organic matter (0.8%), high iron (61 ppm) and high aluminum (124 ppm). Ten individual plants were sampled from each plant species, with a distance of at least 10 meters between each pair of sampled plants, and both fine and woody roots were collected. Samples were kept on ice during travel to the laboratory where they were processed the same day. Root samples were rinsed thoroughly to remove soil from the surface, cut into 10–20 mm lengths using surface sterilized scissors, then surface disinfected with sequential washes of 95% ethanol for 30 s, 0.5% NaOCl for 2 min and 70% ethanol for 2 min. After several rinses with sterile water, root samples were dried and cut into 5 mm pieces using a surface sterilized scalpel then plated on acidified malt extract agar (AMEA, 1.5 ml 85% lactic acid per liter of 2% malt extract agar). Plates were incubated at room temperature with 12 h light and 12 h dark cycles. Fungal cultures were transferred to fresh AMEA and purified by sub-culturing from emergent hyphal tips.

### Morphological study and growth rates

For colony growth rate measurements, isolates PP16K26, PP16K33A, PP16K77A, PP16P16A, PP16P25, and PP16P31 were grown on 2% MEA (BD Difco, Maryland) and 2% water agar (WA) under 12 hr light/12hr dark incubated at 25 C and 30 C with three replicates, and 24 hr dark incubated at 30 C with three replicates. Colony diameter was measured after 14 d. For colony morphology, Cornmeal agar (CMA; BD Difco, Maryland), Czapek’s media (CZM; Wang et al. 2014 BD Difco, Maryland), and potato dextrose agar (PDA; BD Difco, Maryland) were used; while filtered ground pine needle agar (FPNA; Luchi et al. 2007) was used in attempts to induce sporulation; and potato dextrose agar with lecithin (L-PDA; Wang et al. 2014) was used for mating experiments. Cultures were incubated at 25 C in the dark with three replicates and were checked weekly for six mo for mating and sporulation experiments. Capitalized color names used in colony descriptions follow Ridgway (1912).

### DNA extraction, amplification and sequencing

Genomic DNA was extracted from fungal mycelium using the DNeasy PowerSoil isolation kit (Qiagen, Maryland) following the manufacturer’s instructions. PCR was performed with Taq 2X Master Mix (New England BioLabs, Maine), following the manufacturer’s instructions. Primers used were ITS1 and ITS4 for the internal transcribed spacers (ITS1-5.8S-ITS2 = ITS) region, NS1 and NS4 for partial nuc 18S rRNA genes (18S) (White et al. 1990), ITS1 and LR5 for the D1/D2 region of the nuc 28S rRNA genes (28S) (Rehner and Samuels 1995), and fRPB2-5f and fRPB2-7cR (Liu et al. 1999) for the largest subunit of RNA polymerase II (*RPB*2) gene, and Act-1 and Act-4r (Voight and Wöstemeyer 2000) for actin gene (*ACT*). PCR conditions for the ITS, 18S and the 28S consisted of an initial denaturation step at 95 C for 2 min, 35 cycles of 95 C for 45 s, 54 C for 45 s, 72 C for 1.5 min, and a final extension at 72 C for 5 min (Walsh et al. 2018). For *RPB*2 the PCR conditions included an initial denaturation step at 95 C for 2 min, 35 cycles of 95 C for 60 s, 55 C for 2 min, 72 C for 2 min, and a final extension at 72 C for 10 min (Liu et al. 1999). For *ACT* the PCR conditions included an initial denaturation step at 95 C for 2 min, 38 cycles of 94 C for 60 s, 55 C for 60s, 72 C for 60 s, and a final extension at 72 C for 10 min (Wang et al. 2013). PCR products were purified with ExoSAP-IT (Affymetrix, California) following the manufacturer’s instructions and sequenced by Genscript Inc. (Piscataway, NJ) with the same primers used for PCR.

### Sequence alignment and phylogenetic analyses

Seven representative isolates of the new genus *Pygmaeomyces* were included in the phylogenetic analyses along with reference sequences for other Mucoromycotina species (TABLE 1). Sequences were aligned with MUSCLE 4 (Edgar 2004) with a gap opening of −400 and 0 gap extension penalties, and then manually adjusted. The 18S alignment includes the seven new sequences and 21 reference sequences of Mucoromycota (FIG. 1). The purpose of this 18S analysis is to find the phylogenetic position of these new fungal isolates in Mucoromycota. The 3-locus (18S, 28S, *RPB*2) dataset includes the seven new sequences and seven reference sequences of Mucoromycota (FIG. 2). Genealogical concordance was evaluated with the nonparametric Templeton Wilcoxon signed-rank test in PAUP*4.0b10 (Swofford 2002), with 95% bootstrap consensus trees as constraints. No significant conflicts were found between the 18S, 28S, and *RPB*2 gene datasets, so we constructed the combined phylogenetic tree. The individual gene phylogenies from the 3-locus dataset also were constructed (FIGS. 3-5). The ITS alignment includes the seven new sequences and seven environmental sequences from GenBank that have high sequence similarities to them, and four other reference sequences (FIG. 6). The *ACT* alignment includes the seven new sequences and eight reference sequences of Mucoromycota (SUPPLEMENTARY FIG. 1). The variation of taxon sampling in these datasets is due to the difference in sequence data availability. Maximum likelihood (ML) trees were generated with MEGA 6 (Tamura et al. 2013). Models with the lowest BIC scores (Bayesian Information Criterion) were considered to describe the substitution pattern the best. Tamura-Nei 93 was the best model for both 18S datasets, 28S, *ACT* and the 3-gene dataset, while Tamura 3-parameter was the best model for the ITS dataset, and Kimura 2-parameter was the best model for *RPB2*. Initial tree(s) for the heuristic search were obtained automatically by applying Neighbor-Joining and BioNJ algorithms to a matrix of pairwise distances estimated using the Maximum Composite Likelihood approach, and then selecting the topology with superior log likelihood value. A discrete Gamma distribution was used to model evolutionary rate differences among sites. Bootstrap was computed for 500 replications. All positions with less than 95% site coverage were eliminated. Alignments are deposited in TreeBASE (study ID 26509). For the ITS and 28S alignments, Estimates of Evolutionary Divergence analyses were performed. Analyses were conducted on MEGA 6 (Tamura et al. 2013) using the Maximum Composite Likelihood model, and a Gamma distribution was used to model evolutionary rate differences among sites (Tamura et al. 2013). All positions with gaps and missing data were eliminated.

**Figure 1.**
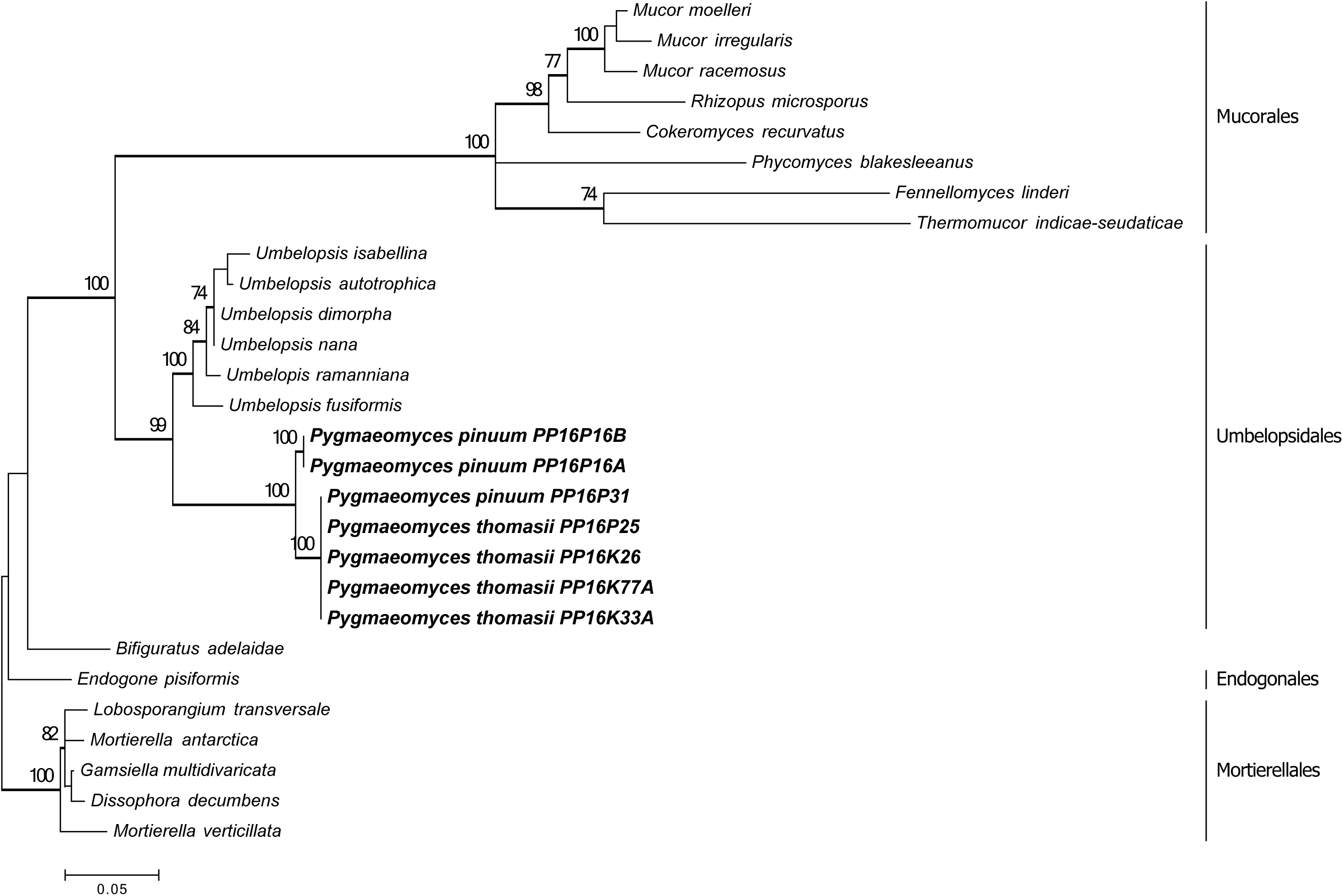
Maximum likelihood phylogenetic tree inferred from 18S gene sequences of *Pygmaeomyces* and 21 reference species of Mucoromycota. Bootstraps higher than 70% have thickened branches. Bar represents substitutions per site.

**Figure 2.**
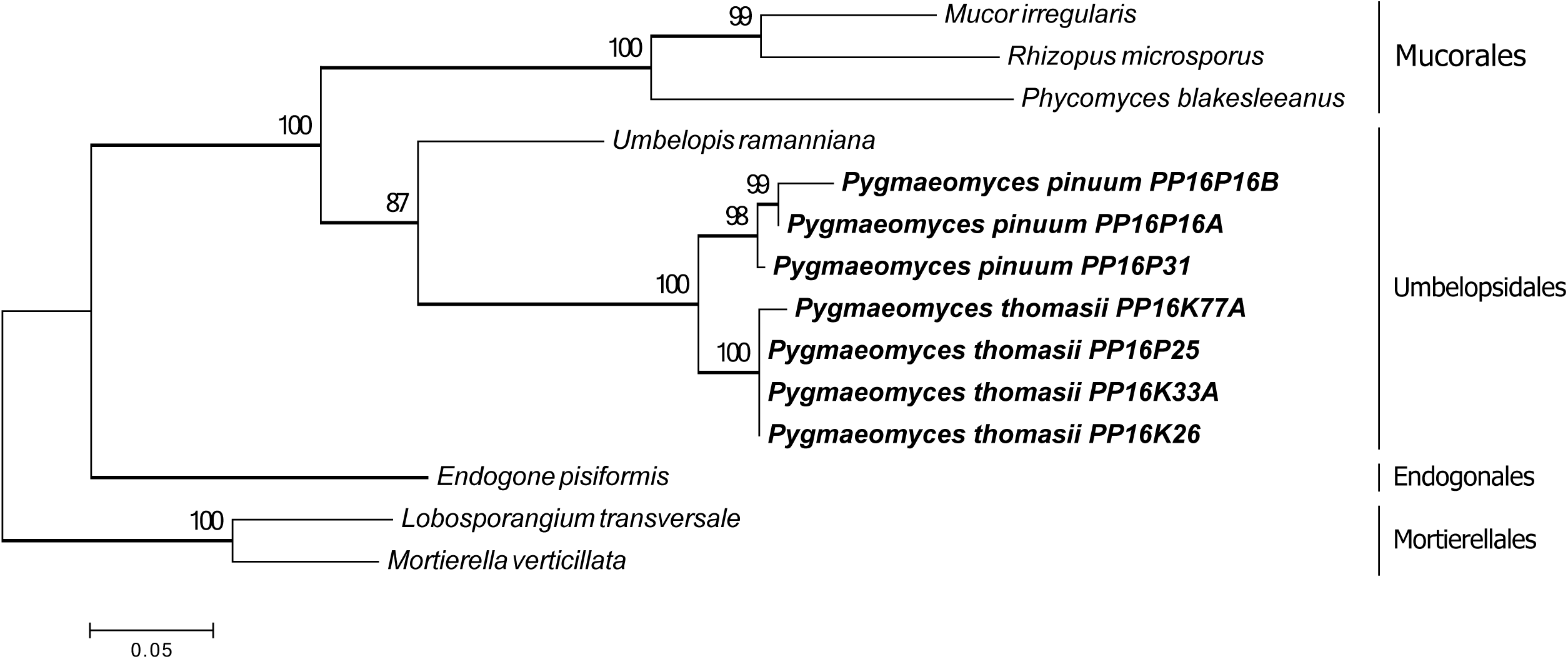
Maximum likelihood phylogenetic tree inferred from combined 18S, 28S and *RPB2* gene sequences of *Pygmaeomyces* and seven reference species. Bootstraps higher than 70% have thickened branches. Bar represents substitutions per site.

**Figure 3.**
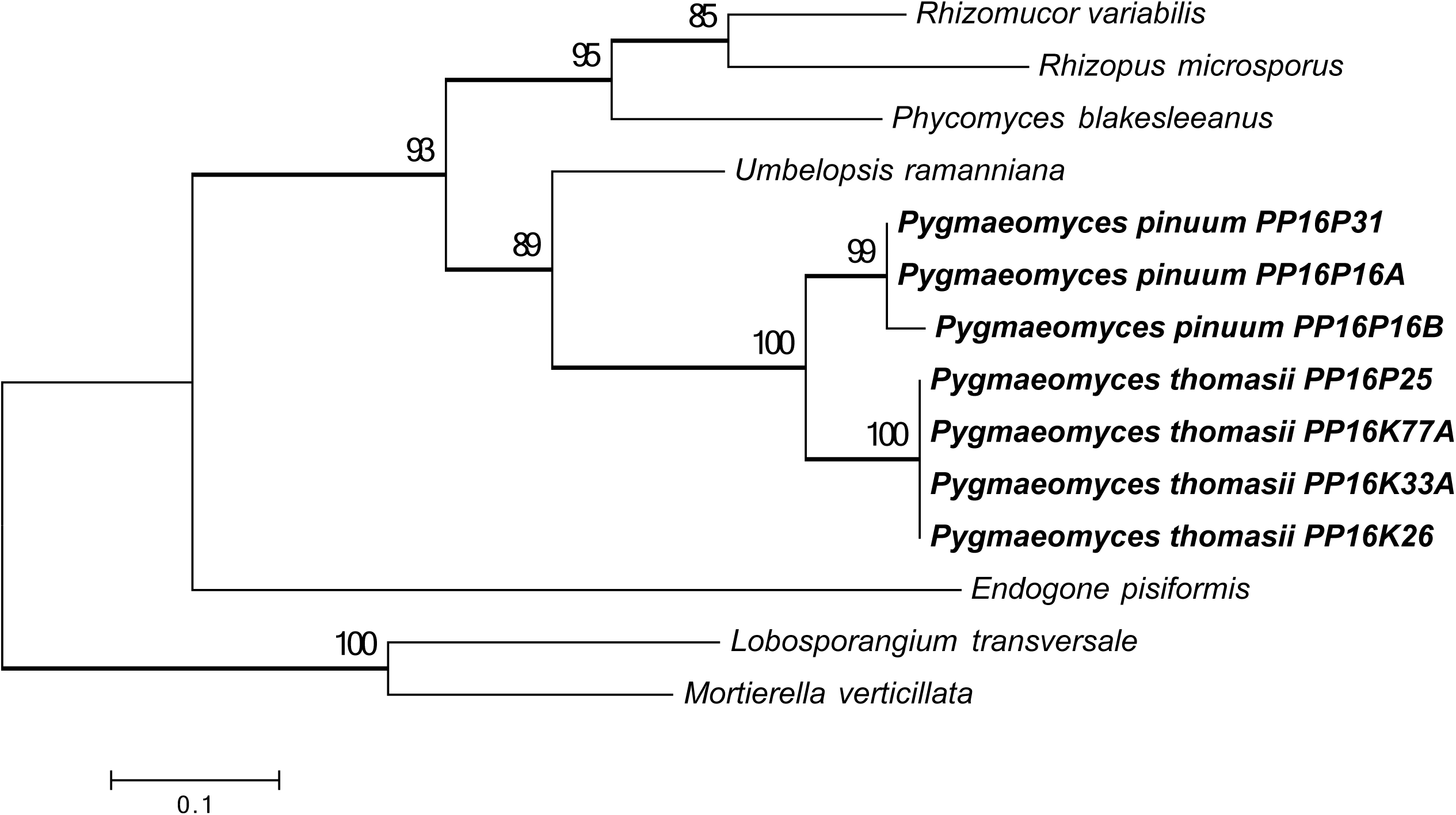
Maximum likelihood phylogenetic tree inferred from *RPB2* gene sequences of *Pygmaeomyces* and seven reference species. Bootstraps higher than 70% have thickened branches. Bar represents substitutions per site.

**Figure 4.**
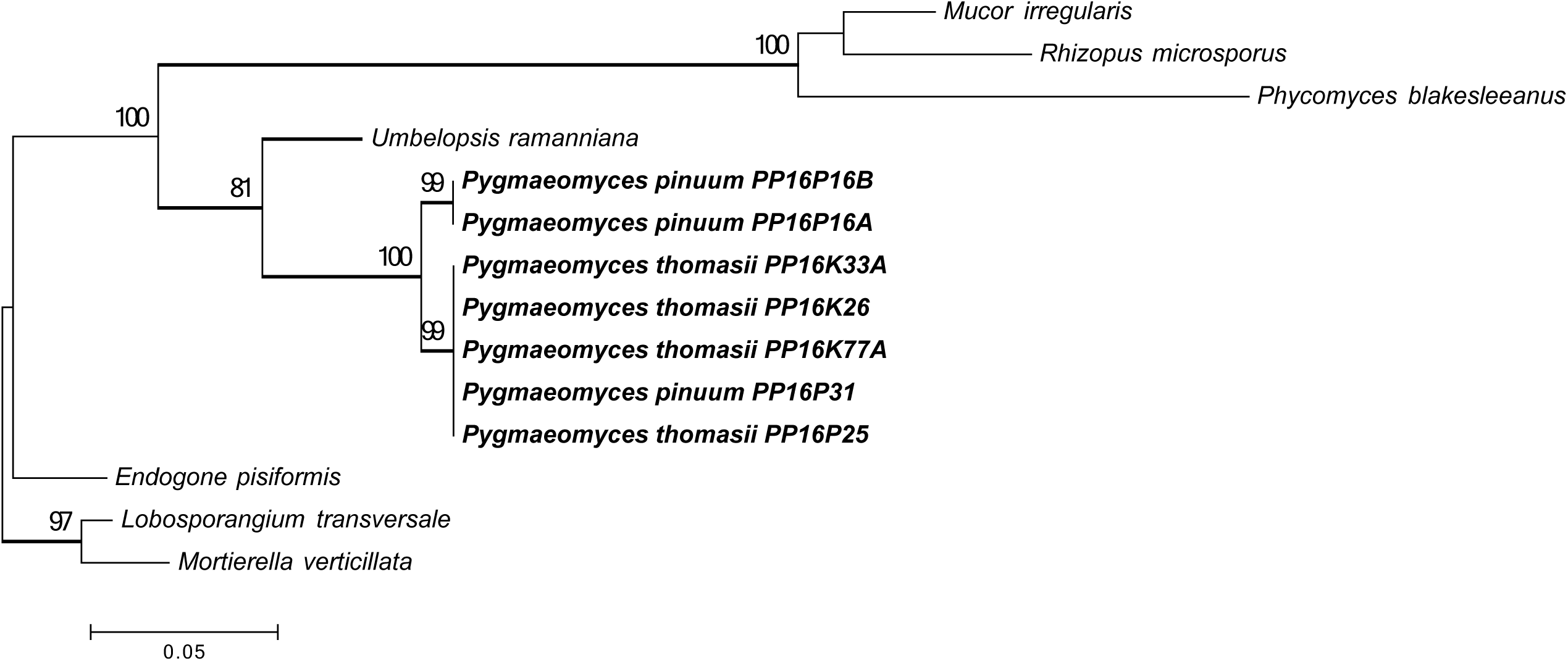
Maximum likelihood phylogenetic tree inferred from the 18S sequences of *Pygmaeomyces* and seven reference species. Bootstraps higher than 70% have thickened branches. Bar represents substitutions per site.

**Figure 5.**
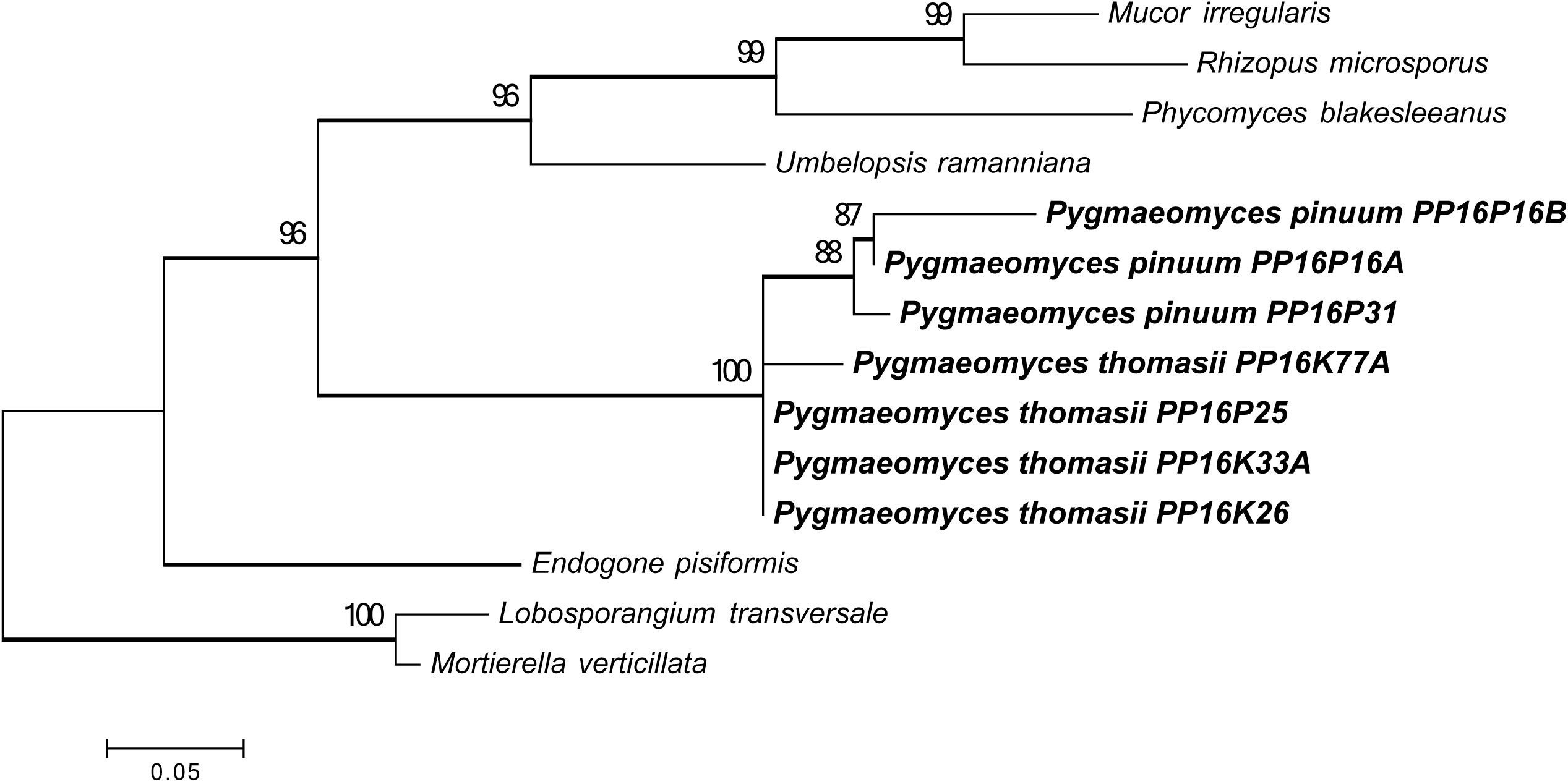
Maximum likelihood phylogenetic tree inferred from the 28S gene sequences of *Pygmaeomyces* and seven reference species. Bootstraps higher than 70% have thickened branches. Bar represents substitutions per site.

**Figure 6.**
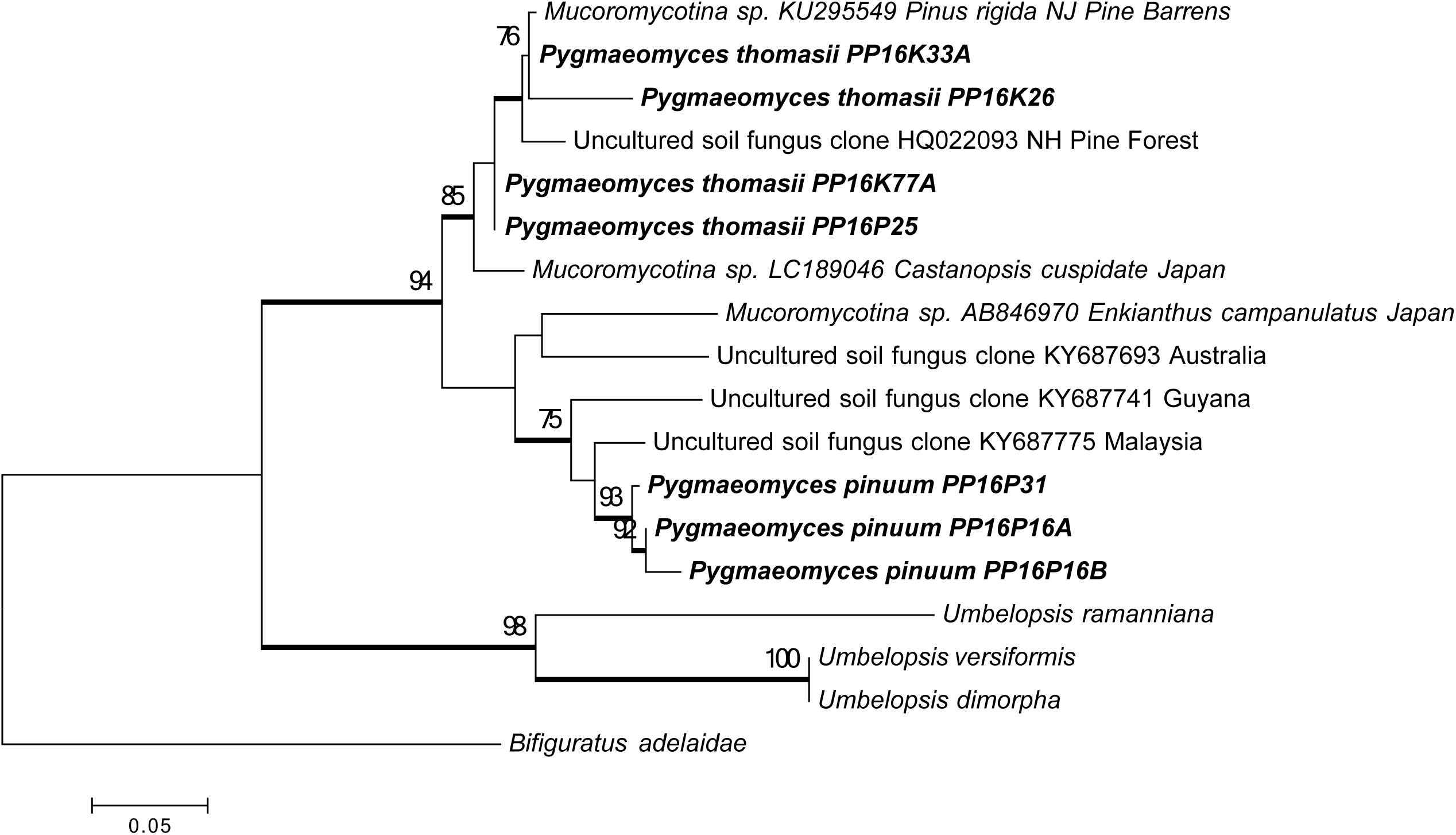
Maximum likelihood phylogenetic tree inferred from the ITS sequences of *Pygmaeomyces* and closely related environmental sequences from GenBank. Bootstraps higher than 70% have thickened branches. Bar represents substitutions per site.

### Plant-fungal interaction experiments

Fungal isolates PP16K26, PP16K77A, and PP16P16A were used in seedling inoculation experiments. Switchgrass (‘Kanlow’) seeds were surface disinfected with 95% ethanol for 30 s, 0.5% NaOCl for 1 min, 70% ethanol for 1 min, rinsed with sterile distilled H_2_O and allowed to germinate in the dark at 25 C for 3 d. Plates of Agargel (Sigma-Aldrich, USA), a medium suitable for plant cell culture, were made following manufacturer’s instructions. The Agargel in the plate was cut in half, and one half of the agargel was removed. On the cut surface of the remaining half Agargel in the plate, three 10 mm × 10 mm × 5 mm plugs from a one-week old fungal culture grown on MEA were placed equidistant from one another. Germinated switchgrass seeds with visible radicle were then placed on the plugs. Sterile MEA plugs were used as a negative control. Cultures were incubated at 25 C under 12 hr light and dark cycle with nine replicates. Root length was recorded at seven days. The same protocol was used for surface sterilized *Kalmia latifolia* seeds. The *Kalmia* root length was recorded at thirty days.

### Microscopy

All images were captured from water mounts by a Nikon DS-Fi1 camera mounted on a Nikon Eclipse 80i compound microscope using the 40× or 60× objectives. Images were measured and analyzed using the Nikon, NIS-Elements D3.0 software.

### Enzyme experiments

The methods of Rice and Currah (2005) were followed to test amylase, gelatinase and lipase activity. Phosphatase, cellulose, and chitinase activity tests followed Pikovskaya (1948), Gupta et al. (2012), and Agrawal et al. (2012), respectively. Cultures of PP16K26, PP16K33A, PP16K77A, PP16P16A, PP16P25, and PP16P31, with three replicates, were grown at room temperature on modified Melin-Norkrans agar (MMN) media plates containing the target macromolecule with or without an indicator (Rice and Currah 2005). Sterile MEA plugs were used as a negative control. Amylase activity was scored after isolates had grown for three wk on plates of MMN containing 2 g/L potato starch, then flooded with iodine solution and decanted after several min to reveal a clear zone around the mycelium in strains positive for this enzyme. Phosphatase activity was scored at seven d on Pikovskaya medium supplemented with 0.025 g/L bromophenol blue (Pikovskaya 1948). Cellulase activity was scored on MMN medium amended with 2g/L carboxymethylcellulose and 0.2 g/L Congo red at five wk (Gupta et al. 2012). Chitinase activity was scored at 14 d on basal chitinase medium amended with 4.5g/L chitin powder and 0.15g/L of bromocresol purple; pH was adjusted to 4.7 before autoclaving (Agrawal et al. 2012). Gelatin medium had 12% gelatin added to MMN media instead of agar, liquefaction after three wk was considered a positive reading for this enzyme. Lipase synthesis on MMN containing 0.1 g/L CaCl_2_ and 10mL/L TWEEN 20 (polyoxyethylene sorbitan monolaurate, Sigma) was determined by the formation of visible crystals beneath the mycelium after 16 wk.

## RESULTS

### Plant-fungal interaction experiments

Switchgrass seedlings inoculated with PP16K26, PP16K77A, and PP16P16A showed no significant differences compared with the control, including the root length and root morphology. *Kalmia* seedlings inoculated with these strains showed no significant differences in root length but the inoculated seedlings exhibited more nodule-like growths along the length of the roots compared to the control (SUPPLEMENTARY FIG. 2).

### Mating experiments and growth on specialty media

For the mating experiment, isolates PP16K26, PP16K33A, PP16K77A, PP16P16A, PP16P25, and PP16P31 were pair-wisely crossed but no zygospore formation was observed. Microchlamydospores were observed from cultures on CMA, CZM, PDA and FPNA but no sporangium was detected.

### Enzyme experiments

PP16K33A, PP16K77A and PP16P25 were negative for phosphatase activity, while PP16K26 scored weakly positive with a faint halo. PP16K26, PP16K77A and PP16P25 were all negative for cellulase activity, while PP16K33A was positive. While PP16P16A, and PP16P31 were weakly positive for cellulase, they were negative for chitinase and phosphatase activity. PP16K26, PP16K33A, PP16K77A and PP16P25 were strongly positive for chitinase. Gelatinase activity was scored positive for all the tested fungi, while amylase activity was negative for all the tested fungi. Lipase synthesis scored a slight reaction for PP16P16A and PP16P25, while all other tested strains scored negative (SUPPLEMENTARY TABLE 1).

### Sequence data and phylogeny

There were 931 characters from the large 18S alignment; 914 from the ITS alignment; 3053 from the three-gene alignment (927 from 18S, 1051 from 28S, and 1075 from *RPB*2); and 1051 from *ACT*. Maximum likelihood trees based on these datasets are shown in FIGS. 1-6 and Supplementary FIG. 1. All phylogenies supported that the new isolates formed a well-supported, monophyletic, distinct clade in Mucoromycotina, and we named it a new genus *Pygmaeomyces*. All except for the 28S tree showed that *Pygmaeomyces* was most closely related to *Umbelopsis*. All except for the 18S trees recognized two groups among the new isolates. PP16K26, PP16K33A, PP16K77A, and PP16P25 formed a well-supported group, described below as *Pygmaeomyces thomasii* while isolates PP16P16A, PP16P16B and PP16P31 formed another, described as *Pygmaeomyces pinuum*. Within-group phylogenetic relationships showed incongruity among different single-gene trees, indicating the occurrence of recombination, which helped delimiting the species boundaries based on the genealogical concordance phylogenetic species recognition (Taylor et al. 2000). Evolutionary Divergence analyses of ITS sequences (SUPPLEMENTARY TABLE 2) showed a range of 84-91% similarities between the isolates of *Pygmaeomyces thomasii* and *P. pinuum*, 62-71% between *Pygmaeomyces* and *Umbelopsis* species, and 58-60% between *Pygmaeomyces* and Endogonales. Evolutionary Divergence analyses of the 28S sequences (SUPPLEMENTARY TABLE 3) showed 73-83% similarities between *Pygmaeomyces* and Umbelopsidaceae, 63-75% between *Pygmaeomyces* and Endogonaceae, and 45-76% between *Pygmaeomyces* and Mucoraceae. The percent similarities between a number of closely related families in Mucoromycotina based on the 28S sequences are as follows: between Densosporaceae and Endogonaceae 79-81%, between Pilobolaceae and Rhizopodaceae 87-92%, Lichtheimiaceae and Mucoraceae 57-84%, Cunninghamellaceae and Mucoraceae 51-68%, Lichtheimiaceae and Cunninghamellaceae 51-68%, Radiomycetaceae and Phycomycetaceae 60-71%, Saksenaeaceae and Mucoraceae 54-79%, Choanephraceae and Mucoraceae 74-91%, and Syncephalastraceae and Lichtheimiaceae 50-65%. The 28S sequence similarities between the *Pygmaeomyces* clade and closely related families are: Umbelopsidaceae 73-83%, Endogonaceae 63-75%, and Mucoraceae 45-76%. Based on the molecular phylogenetic analyses, divergence analyses, morphological characters and their ecological features, two new species, a new genus and a new family are proposed.

## TAXONOMY

**Pygmaeomycetaceae** E. Walsh & N. Zhang, fam. nov.

MycoBank: MB832250

*Typification*: *Pygmaeomyces* E. Walsh & N. Zhang

*Diagnosis*: Pygmaeomycetaceae is erected here to apply to all descendants of the node defined in the combined 18S, 28S, and *RPB2* phylogeny (FIG. 2) as the terminal clade containing the genus *Pygmaeomyces.* Phylogenetic analyses place this as a sister group to Umbelopsidaceae in Umbelopsidales, consistent with family status. Distinguished from other families in the Mucoromycotina by producing hyaline microchlamydospores.

*Description*: Associated with roots of plants in acidic soils. Subglobose vesicles formed from coenocytic hyaline hyphae. Microchlamydospores hyaline, globose to subglobose. Sporangia not observed. Sexual reproduction unknown.

***Pygmaeomyces*** E. Walsh & N. Zhang, gen. nov.

MycoBank: MB832252

*Typification: Pygmaeomyces thomasii* E. Walsh & N. Zhang

*Etymology*: “pygmaeus” means Pygmy, referring to the Pygmy pine plains ecosystem where the fungi were discovered.

*Diagnosis*: In addition to the phylogenetic distinctions (FIGS. 1-6), *Pygmaeomyces* differs from *Umbelopsis* by the lack of sporangiophores and sporangiospores, and the lack of reddish or ochraceous pigmentation.

*Description:* Colonies on MEA lightly pigmented, mucoid textured surface thick and light brown, sparse aerial hyphae if present. Colonies on WA light brown, mucoid textured surface with sparse aerial hyphae if present. Subglobose vesicles formed from coenocytic hyaline hyphae. Microchlamydospores hyaline, globose to subglobose. Sporangia and sexual reproduction unknown.

*Habitat*: Associated with roots of plants in acidic soils.

*Known distribution*: New Jersey, USA.

***Pygmaeomyces thomasii*** E. Walsh & N. Zhang, sp. nov. (FIG. 7B, E-J)

**Figure 7.**
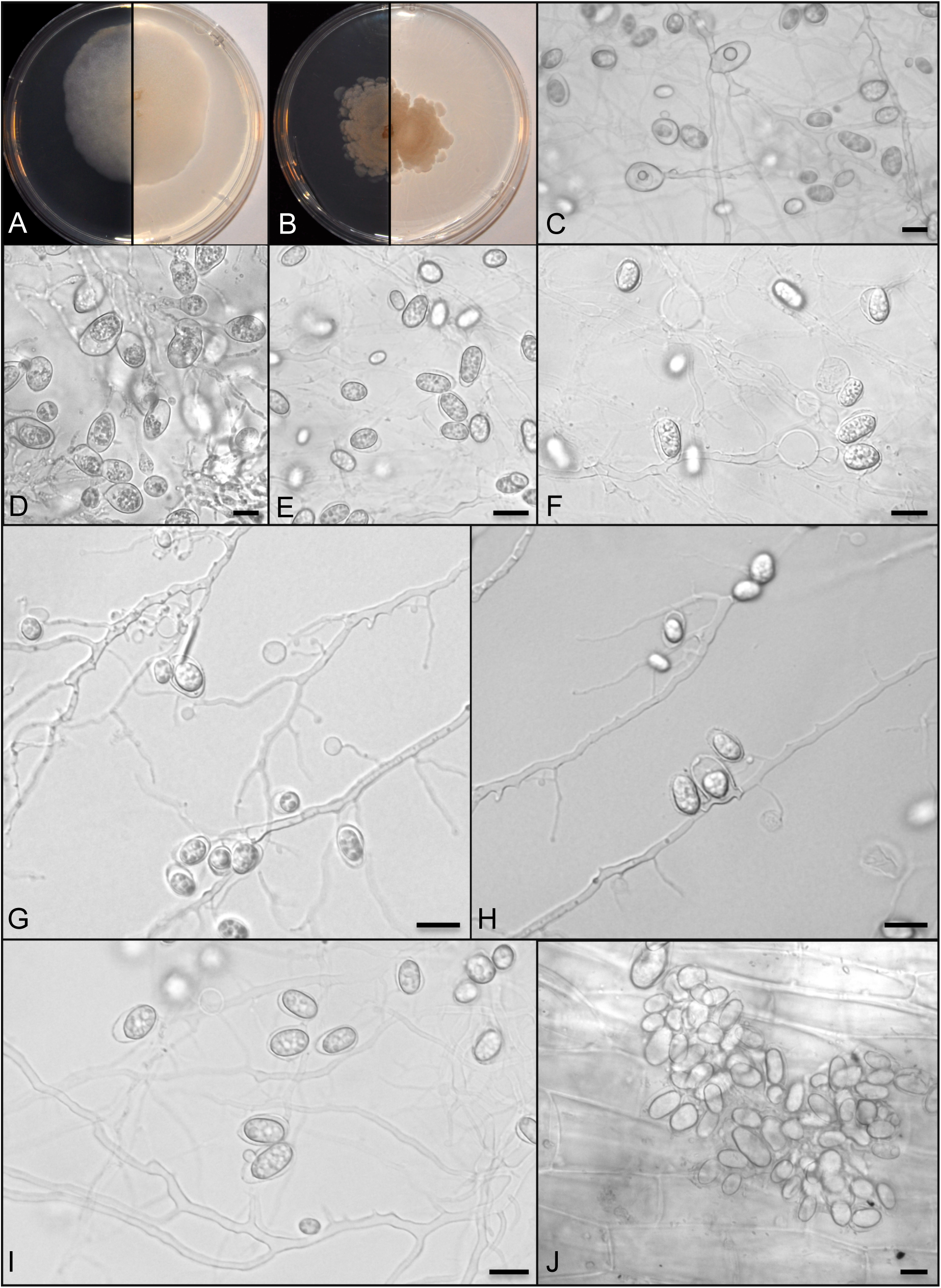
Cultures on MEA+LA 60mm plates A. *Pygmaeomyces pinuum*. B. *Pygmaeomyces thomasii*, Microchlamydospores C-D, From *Pygmaeomyces pinuum* ex-type culture. E–I. From *Pygmaeomyces thomasii* ex-type culture, J. Switchgrass seedling roots inoculated with *Pygmaeomyces thomasii* ex-type culture. All bars = 10 μm.

MycoBank: MB832253

*Typification:* USA. NEW JERSEY: West Pygmy Pine Plains, Little Egg Harbor, 39.708783, −74.372567, 26 m alt, from roots of apparently healthy *Pinus rigida* and *Kalmia latifolia* in a pygmy pine forest, 22 Jun 2016, *E. Walsh & N. Zhang PP16K26* (**holotype** RUTPP-PP16K26). Ex-type culture (CBS 146528). GenBank: ITS = MN017028; *RPB*2= MN486053.

*Etymology:* The epithet honors the first author’s late father, Thomas A. Walsh, who encouraged her in her scientific and research endeavors.

*Description:* Colonies on MEA 26 mm diam on average with standard deviation (SD) of 5.3 after 7 d in the light at 25 C, Fawn Color, mycelium with a Cinnamon margin, sparse aerial hyphae light brown, reverse pigmented, Cinnamon. Colonies on MEA 24 mm diam on average with SD of 2 after 7 d in the dark at 25 C, and 17.3 mm diam on average with SD of 2.51 after 7 d in the dark at 30 C. Colonies on WA 7 mm diam on average with SD of 0 after 7 d in the dark at 25 C, Wood Brown, aerial hyphae sparse, reverse pigmented, Avellaneous. Colonies on WA 6.7 mm diam on average with SD of 0.6 after 7 d in the dark at 25 C, and 7 mm diam on average with SD of 0 after 7 d in the dark at 30 C. Coenocytic hyaline hyphae becoming septate to form subglobose to globose vesicles. No sporangia were observed. Microchlamydospores hyaline, globose to subglobose, 6.24–15 × 4.11–8 μm (n = 100, mean 9.92 × 6.05 μm, s.e. 1.59, 0.9). Zygospores not observed.

*Other material examined:* USA. NEW JERSEY: same location as ex-type culture, from roots of apparently healthy *Pinus rigida* and *Kalmia latifolia* in the pygmy pine plains, 22 Jun 2016, *E. Walsh & N. Zhang PP16K33A, PP16K77A* and *PP16P25*.

***Pygmaeomyces pinuum*** E. Walsh & N. Zhang, sp. nov. (FIG. 7A, C-D)

MycoBank: MB832254

*Typification:* USA. NEW JERSEY: West Pygmy Pine Plains, Little Egg Harbor, 39.708783, −74.372567, 26 m alt, from roots of apparently healthy *Pinus rigida* in a subalpine forest, 22 Jun 2016, *E. Walsh & N. Zhang PP16P16A* (**holotype** RUTPP-PP16P16A). Ex-type culture CBS 146529). GenBank: ITS = MN017032; *RPB2*= MN486057.

*Etymology:* The epithet refers to *Pinus*, the host plant of the fungi.

*Description:* Colonies on MEA 34.6 mm diam on average with SD of 1.53 after 7 d in the light at 25 C, dense velvet-like, Vinaceous Buff, sparse aerial hyphae, reverse pigmented, Colonial Buff. Colonies on MEA 30.3 mm diam on average with SD of 0.5 after 7 d in the dark at 25 C, and 44 mm diam on average with SD of 1.7 after 7 d in the dark at 30 C. Colonies on WA reaching 6.7 mm diam on average with SD of 0.6 after 7 d in the light at 25 C, sparse Avellaneous mycelium with a Wood Brown margin, reverse pigmented, Avellaneous. Colonies on WA 6 mm diam on average with SD of 0 after 7 d in the dark at 25 C, and 6 mm diam on average with SD of 0 after 7 d in the dark at 30 C. Coenocytic hyaline hyphae became septate to form subglobose vesicles. No sporangia were observed. Microchlamydospores hyaline, globose to subglobose, 6.06–14.69 × 2.67–10.13μm (n = 100, mean 9.89 × 6.74 μm, s.e. 1.59, 1.35). Zygospores not observed.

*Other material examined:* USA. NEW JERSEY: same location as ex-type culture, from roots of apparently healthy *Pinus rigida* in the pygmy pine plains, 22 Jun 2016, *E. Walsh & N. Zhang PP16P16B*, and *PP16P31.*

*Notes:* In addition to the phylogenetic distinctions (FIG. 2), *Pygmaeomyces pinuum* differs from *P. thomasii* by dense velvet like aerial hyphae.

## DISCUSSION

Zygomycete fungi have long been recognized to be non-monophyletic (Hibbett et al. 2007). Based on phylogenomic analyses of 46 taxa including 25 zygomycetes, Spatafora et al. (2016) recently reclassified the zygomycete fungi into two phyla, Mucoromycota and Zoopagomycota. Mucoromycota is comprised of three subphyla: Glomeromycotina, Mortierellomycotina, and Mucoromycotina. The new taxa described in this paper belong to the order Umbelopsidales in Mucoromycotina. Endogonales and Mucorales are the other two orders in Mucoromycotina. Typically, fungi in Mucoromycotina have fast growing coenocytic hyphae, produce sporangia, sporangioles or chlamydospores, and are mycorrhizal, root endophytes or saprobes (Hibbett et al. 2007, Benny et al. 2014, Spatafora et al. 2016), which match well with the characteristics of *Pygmaeomyces.*

The phylogenetic analyses in this study indicated that *Pygmaeomyces* formed a well-supported, distinct clade in Mucoromycotina and its closest relative is *Umbelopsis. Umbelopsis* species usually produce sporangia while *Pygmaeomyces* species only produce microchlamydospores. Moreover, *Pygmaeomyces* species have 71 % or less ITS sequence similarities to *Umbelopsis* species and any other described taxa with accessible ITS sequences from GenBank. The 71% ITS sequence similarity is significantly lower than the commonly observed 97-99.6% ITS similarities at species level of filamentous fungi (Walsh et al. 2014, 2015; Vu et al. 2019), which justifies the novelty status of the *Pygmaeomyces* species. The new family status of Pygmaeomycetaceae was based on the phylogeny and the 28S sequence distance/similarity comparison. These numbers fall into the 28S sequence similarity range at family level in Mucoromycotina listed above; therefore, we propose a new family, Pygmaeomycetaceae in the order Umbelopsidales (Spatafora et al. 2016, Tedersoo et al. 2018). Tedersoo et al (2017) used 80% ITS sequence similarity for the family or order distinction. Vu et al. (2019) suggested a 96.2% 28S sequence threshold for family level in filamentous fungal identification, which is significantly higher than the observed percent similarities in the established families in Mucoromycotina. Fungi are highly diverse and genetically variable, and the discrepancy observed here indicates that a universal sequence similarity threshold does not apply for all fungal lineages. To perform robust and meaningful taxon recognition, we should consider all available information, including phylogenetic analyses, sequence similarity, phenotypic, ecological, and other characters if available.

The GenBank BLAST results and phylogenetic analyses indicated that fungi in the *Pygmaeomyces* clade likely have a wide distribution. Fourteen environmental ITS sequences in GenBank had 90-94 % identities with that of *Pygmaeomyces pinuum*, for example, KY687775 from tropical rainforest soil in Malaysia, KY687741 from tropical rainforest soil in Guyana, and AB846970 from roots of *Enkianthus campanulatus* in Japan. Fourteen ITS sequences in GenBank had 90-98 % identities with that of *Pygmaeomyces thomasii*, for example, KU295549 from roots of *Pinus rigida* in the New Jersey Pine Barrens, LC189046 from roots of *Castanopsis cuspidata* in Japan and HQ022093 from temperate forest soil in USA. Most of these sequences were from either acidic soils or the roots of Ericaceae or conifers that usually grow in acidic soils, which is similar to the edaphic condition of the New Jersey Pine Plains. Interestingly, some of these *Pygmaeomyces*-similar sequences (KY687741, KY687775, KY687693) were generated from a culture-independent soil fungal DNA survey and they have been named as Clade GS23 (Tedersoo et al. 2017). Their phylogenetic analysis similarly placed Clade GS23 as a monophyletic branch to Umbelopsidaceae (Tedersoo et al. 2017). Clade GS23 contains environmental samples from tropical rain forests with very low soil pH from Australia, Guyana, and Malaysia. The affinity for low pH soils and a global distribution pattern was also found in the genera *Acidomelania* and *Barrenia*, root associated fungi frequently isolated from the acidic pine barrens (Walsh et al. 2014, Walsh et al. 2015).

The functions of *Pygmaeomyces* have not been fully understood but the fact that they were isolated from apparently healthy plants, and no negative impact observed in the plant-fungal interaction experiments indicate that they are not pathogens. Moreover, their affinity to plants in acidic soils suggests that these fungi may have adapted to such low pH environment or may have helped their host plants to adapt to it. All of the *Pygmaeomyces* species were able to produce chitinases and gelatinases and one isolate was able to degrade cellulose. These results suggest that these enzymes potentially allow them to degrade various substrates, such as plants, fungi, insects or other animals. Bååth and Söderström (1980) found that *Mortierella* and *Mucor* isolates from an acidic coniferous forest also produced chitinases. The ability to degrade chitin by these fungi allows for the mobilization of nitrogen and may contribute to the success of their host plants in the acidic, oligotrophic environments.

The novel fungal lineages identified based on environmental sequencing, such as Clade GS23 (Tedersoo et al. 2017) may be called “dark taxa” or “dark matter fungi”-undescribed fungal taxa only known from sequence data without a physical specimen and lacking a resolved taxonomic identity (Grossart et al. 2016, Ryberg and Nilsson 2018). These “dark taxa” are believed to be a combination of undescribed novel lineages and described taxa without sequencing data (Nagy et al. 2011). Here we linked the “dark taxon” GS23 with the fungal cultures isolated from the plant roots in the pine plains. Another example of bridging the gap is the isolation and description of *Bifiguratus*, the sequence of which was known as UCL7_006587 (Torres-Cruz et al. 2017). A concern about the sequence-only analyses is that they tend to inflate the number of taxon names (or MOTU) (Clare et al. 2016, Kunin et al. 2010, Ryberg and Nilsson 2018). We observed that sequence-only analyses also have the tendency to over-estimate the taxonomic level. GS23, along with a number of other lineages recognized in the soil survey by Tedersoo et al. (2017) was believed to have at least class level distinction from other fungi. However, our culture-based analyses placed them at family level. The current problems in environmental sequencing, such as short sequence length, low quality and chimera may explain the observed difference between the sequence-only and culture-based taxonomic analyses.

In summary, we described a new family, a new genus and two new species of root-colonizing fungi associated with plants living in acidic soils. The phylogenetic and taxonomic work and the plant–fungal interaction results reported here will aid future ecological and evolutionary studies on root-associated fungi, and will help elucidating the function of these root symbionts in the understudied pygmy pine plains. The work presented here also provides an example on linking the culture-based and sequence-only fungal identification.

## ACKNOWLEDGMENTS

The research was supported by the National Science Foundation (DEB 1452971) and the Rutgers University Alberts Research Awards in Biodiversity. The first author would like to thank her father, T.A. Walsh, who enjoyed hearing tales of her collecting trips and about the progress of her research, experiments, and papers. And as an English Literature teacher fond of puns, he found a way to incorporate fungi into the everyday; he became a man of morels and was overall a truly fun guy.

## LEGENDS and FOOTNOTES

Table 1. Species name, isolate number, host location and GenBank accession numbers of the fungi used in this study.

Suppl. Table 1. Enzyme tests results

Suppl. Table 2. Estimates of Evolutionary Divergence inferred from the ITS sequence data set.

Suppl. Table 3. Estimates of Evolutionary Divergence inferred from the 28S sequence data set.

Suppl. Figure 1. Maximum likelihood phylogenetic tree inferred from *ACT* gene sequences. Bootstraps higher than 70% have thickened branches. Bar represents substitutions per site.

Suppl. Figure 2. *Kalmia latifolia* seedlings fourteen days after inoculation on 60 mm agargel plates. A. Control *Kalmia latifolia* seedlings B. *Pygmaeomyces thomasii* isolate PP16K77A. C. *Pygmaeomyces pinuum* holotype isolate PP16P16A.

