## Supplementary material for "Pygmaeomycetaceae, a new root associated family in Mucoromycotina from the pygmy pine plains": TABLE 1

Table 1. Species name, isolate number, host, location and GenBank accession numbers of the fungi used in this study.

| Species | Isolate number ^a^ | Host | Location | ITS | 28S | 18S | *RPB2* | *ACT* |
| --- | --- | --- | --- | --- | --- | --- | --- | --- |
| *Bifiguratus adelaidae* | NRRL66559 | leaf tissue of *Leucobryum* sp. | Arizona, USA | **MN062398** ^b^ | HM123225 | KX372677 |  | **MN313255** |
| *Cokeromyces recurvatus* | CBS158.50 | dung of rabbit | Illinois, USA |  |  | AY635843 |  |  |
| *Dissophora decumbens* | CBS592.88 | ground-up *Quercus* and *Acer* leaves | Rhode Island, USA |  |  | HQ667440 |  |  |
| *Endogone pisiformis* | AFTOL539 |  |  | AY997046 | DQ273811 | DQ322628 | DQ302776 | AB609182 |
| *Fennellomyces linderi* | CBS158.54 | poplin | Florida, USA |  |  | JX629074 |  |  |
| *Gamsiella multivaricata* | CBS227.78 | decaying stump | Moscow, Russia |  |  | HQ667475 |  |  |
| *Gigaspora margarita* |  | *Trifolium repens* |  |  |  |  |  | AJ567829 |
| *Lobosporangium transversale* | NRRL3116 |  |  |  | HQ667404 | HQ667488 | XM_022027218 | AJ287156 |
| *Mortierella antarctica* | CBS609.70 | soil near glacier | Antarctica |  |  | HQ667503 |  |  |
| *Mortierella verticillata* | NRRL6337 =AFTOL141 | sandy forest soil | United Kingdom | AY997063 | DQ273794 | AF157145 | DQ302784 | AJ287170 |
| *Mucor irregularis* | CBS103.93 | wrist/back of hand | Jiangsu, China | JN206150 | JX976212 | HM623315 | JX976279 |  |
| *Mucor moelleri* | CBS444.65 | soil | Wyoming, USA |  |  | EU736298 |  |  |
| *Mucor racemosus* | CBS260.68 |  | Basel, Switzerland |  |  | JF723672.2 |  |  |
| Mucoromycotina sp. |  | roots of *Castanopsis cuspidate* | Japan | LC189046 |  |  |  |  |
| Mucoromycotina sp. |  | roots of *Ekianthus campanulatus* | Japan | AB846970 |  |  |  |  |
| Mucoromycotina sp. | WSF14P94 | roots of *Pinus rigida* | New Jersey, USA | KU295549 |  |  |  |  |
| *Phycomyces blakesleeanus* | AFTOL184 |  |  | AY997071 | DQ273800 | AY635837 | DQ302789 |  |
| *Pygmaeomyces pinuum* | PP16P16A= CBS146529 | roots of *Pinus rigida* | New Jersey, USA | **MN017032** | **MN017097** | **MN017039** | **MN486057** | **MN313252** |
| *Pygmaeomyces pinuum* | PP16P16B | roots of *Pinus rigida* | New Jersey, USA | **MN017033** | **MN017098** | **MN017040** | **MN486058** | **MN313253** |
| *Pygmaeomyces pinuum* | PP16P31 | roots of *Pinus rigida* | New Jersey, USA | **MN017034** | **MN017099** | **MN017041** | **MN486059** | **MN313254** |
| *Pygmaeomyces thomasii* | PP16K26= CBS146528 | roots of *Kalmia latifolia* | New Jersey, USA | **MN017028** | **MN017093** | **MN017035** | **MN486053** | **MN313248** |
| *Pygmaeomyces thomasii* | PP16K33A | roots of *Kalmia latifolia* | New Jersey, USA | **MN017029** | **MN017094** | **MN017036** | **MN486054** | **MN313249** |
| *Pygmaeomyces thomasii* | PP16K77A | roots of *Kalmia latifolia* | New Jersey, USA | **MN017030** | **MN017095** | **MN017037** | **MN486055** | **MN313250** |
| *Pygmaeomyces thomasii* | PP16P25 | roots of *Pinus rigida* | New Jersey, USA | **MN017031** | **MN017096** | **MN017038** | **MN486056** | **MN313251** |
| *Rhizopus microsporus* | CBS699.68 = ATCC52813 | soil | Ukraine | JN206364 | FN182233 | FN182238 | XM_023614055 |  |
| *Thermomucor indicae-sedudaticae* | CBS104.75 | municipal compost | Poona, India |  |  | AF157165 |  |  |
| *Umbelopsis autotrophica* | CBS310.93 | Soil | Oxshott Heath, UK |  |  | KM017660 |  |  |
| *Umbelopsis dimorpha* | CBS110039 | soil | New Zealand | JN206387 | KF727471 | KM017689 |  | KF771833 |
| *Umbelopsis fusiformis* | CBS385.85 | Soil from forest of *Eucalyptus, regnans* | Melbourne, Australia |  |  | KM017683 |  |  |
| *Umbelopsis isabellina* | CBS208.32 | Sandy loam | Victoria, Australia |  |  | KM017681 |  |  |
| *Umbelopsis nana* | CBS730.70  NRRL22420 | Forest soil | The Netherlands |  |  | KM017695 |  |  |
| *Umbelopsis ramanniana* | NRRL5844  =AFTOL144 |  |  | KM017730 | DQ273797 | KM017694 | DQ302787 | KM017713 |
| *Umbelopsis versiformis* | CBS150.81 | *Quercus borealis,* root | Virginia, USA | KC489496 |  |  |  | KF771851 |
| Uncultured fungus clone |  | soil | Malaysia | KY687775 |  |  |  |  |
| Uncultured fungus clone |  | soil | New Hampshire, USA | HQ022093 |  |  |  |  |
| Uncultured fungus clone |  | soil | Australia | KY687693 |  |  |  |  |
| Uncultured fungus clone |  | soil | Guyana | KY687775 |  |  |  |  |

^a^ AFTOL= Assembling the Fungal Tree of Life project; ATCC/MYA = American Type Culture Collection, Manassas, Virginia, USA; CBS = Centraalbureau voor Schimmelcultures, Utrecht, The Netherlands; NRRL/ARS = Agricultural Research Service Culture Collection, Peoria, Illinois, USA.

^b^ Numbers in boldface indicating new sequences from this study.
