## SUPPLEMENTARY TABLE 1 for "Pygmaeomycetaceae, a new root associated family in Mucoromycotina from the pygmy pine plains"

Supplementary Table 1. Enzyme test results

|  | Isolate # | AMYLASE | | | CELLULASE | | | CHITANASE | | | GELATINASE | | | LIPASE | | | PHOSPHATASE | | |
| --- | --- | --- | --- | --- | --- | --- | --- | --- | --- | --- | --- | --- | --- | --- | --- | --- | --- | --- | --- |
|  |  | 1 | 2 | 3 | 1 | 2 | 3 | 1 | 2 | 3 | 1 | 2 | 3 | 1 | 2 | 3 | 1 | 2 | 3 |
| *Acidomelania panicicola* | CBS137156 | - | - | - | +++ | +++ | +++ | - | - | - | ++ | ++ | ++ | - | - | - | + | + | - |
| *Barrenia panicia* | WSF1R37 | + | + | + | ++ | ++ | ++ | + | + | + | - | - | - | - | - | - | + | + | + |
| *Cadophora meredithiae* | BAG2 | - | - | - | + | + | + | - | - | - | +++ | +++ | +++ | ++ | + | - | - | - | - |
| *Cadophora interclivum* | BAG4 | - | - | - | - | - | - | - | - | - | + | + | + | - | - | - | - | - | - |
| *Umbelopsis dimorpha* | PP16-P60 | ++ | ++ | ++ | +++ | +++ | +++ | ++ | ++ | ++ | +++ | +++ | +++ | - | - | - | +++ | +++ | +++ |
| *Umbelopsis ramanniana* | PP16-P33B | +++ | ++ | +++ | +++ | +++ | +++ | +++ | +++ | +++ | +++ | +++ | +++ | - | - | - | +++ | +++ | +++ |
| *Pygmaeomyces pinuum* | P16A | - | - | - | + | + | + | - | - | - | +++ | +++ | +++ | + | + | + | - | - | - |
| *Pygmaeomyces pinuum* | P31 | - | - | - | + | + | + | - | - | - | +++ | +++ | +++ | - | - | - | - | - | - |
| *Pygmaeomyces thomasii* | K26 | - | - | - | - | - | - | + | ++ | ++ | +++ | ++ | ++ | - | - | - | ++ | + | + |
| *Pygmaeomyces thomasii* | K33A | - | - | - | +++ | +++ | +++ | ++ | ++ | +++ | +++ | ++ | ++ | - | - | - | - | - | - |
| *Pygmaeomyces thomasii* | K77A | - | - | - | - | - | - | ++ | +++ | ++ | +++ | +++ | +++ | - | - | - | - | - | - |
| *Pygmaeomyces thomasii* | P25 | - | - | - | - | - | - | ++ | +++ | ++ | +++ | +++ | +++ | + | + | - | - | - | - |
| *Bifiguratus adelaide* | AZ0501 | ++ | ++ | ++ | + | + | + |  |  |  | ++ | ++ | ++ |  |  |  | ++ | ++ | ++ |

- means no reaction, + indicates a slight reaction, ++ means positive, +++ means definitely positive
