## SUPPLEMENTARY TABLE 2 for "Pygmaeomycetaceae, a new root associated family in Mucoromycotina from the pygmy pine plains"

| | | 1 | 2 | 3 | 4 | 5 | 6 | 7 | 8 | 9 | 10 | 11 | 12 | 13 | 14 | 15 | 16 | 17 | 18 |
|---|---|---|---|---|---|---|---|---|---|---|---|---|---|---|---|---|---|---|---|
| 1 | Mucoromycotina sp. WSF14 P94 Pine Barrens | - | | | | | | | | | | | | | | | | | |
| 2 | Mucoromycotina sp. N2-1.1-NN-2016 Japan | 0.04 | | | | | | | | | | | | | | | | | |
| 3 | Mucoromycotina sp. 4 KO-2013 Japan | 0.12 | 0.14 | | | | | | | | | | | | | | | | |
| 4 | Uncultured fungus clone Guyana | 0.12 | 0.12 | 0.14 | | | | | | | | | | | | | | | |
| 5 | Uncultured fungus clone Malaysia | 0.10 | 0.11 | 0.13 | 0.08 | | | | | | | | | | | | | | |
| 6 | Uncultured soil fungus clone NH Pine Forest | 0.02 | 0.04 | 0.13 | 0.10 | 0.09 | | | | | | | | | | | | | |
| 7 | Uncultured fungus clone G23 | 0.12 | 0.13 | 0.14 | 0.15 | 0.11 | 0.14 | | | | | | | | | | | | |
| 8 | Pygmaeomyces thomasii PP16K26 | 0.04 | 0.08 | 0.16 | 0.15 | 0.12 | 0.06 | 0.15 | | | | | | | | | | | |
| 9 | Pygmaeomyces thomasii PP16K33A | 0.00 | 0.04 | 0.12 | 0.11 | 0.09 | 0.02 | 0.11 | 0.04 | | | | | | | | | | |
| 10 | Pygmaeomyces thomasii PP16K77A | 0.02 | 0.03 | 0.12 | 0.11 | 0.10 | 0.03 | 0.11 | 0.06 | 0.01 | | | | | | | | | |
| 11 | Pygmaeomyces thomasii PP16P25 | 0.02 | 0.03 | 0.12 | 0.11 | 0.10 | 0.03 | 0.11 | 0.06 | 0.01 | 0.00 | | | | | | | | |
| 12 | Pygmaeomyces pinuum PP16P16A | 0.10 | 0.10 | 0.12 | 0.08 | 0.04 | 0.09 | 0.10 | 0.14 | 0.10 | 0.09 | 0.09 | | | | | | | |
| 13 | Pygmaeomyces pinuum PP16P16B | 0.12 | 0.12 | 0.14 | 0.10 | 0.06 | 0.11 | 0.12 | 0.16 | 0.12 | 0.11 | 0.11 | 0.01 | | | | | | |
| 14 | Pygmaeomyces pinuum PP16P31 | 0.10 | 0.10 | 0.12 | 0.07 | 0.04 | 0.09 | 0.10 | 0.15 | 0.10 | 0.10 | 0.10 | 0.01 | 0.02 | | | | | |
| 15 | Umbelopsis ramanniana strain NRRL 5844 | 0.35 | 0.34 | 0.36 | 0.37 | 0.34 | 0.35 | 0.35 | 0.38 | 0.34 | 0.34 | 0.34 | 0.34 | 0.37 | 0.35 | | | | |
| 16 | Umbelopsis versiformis strain CBS 150.81 | 0.30 | 0.28 | 0.33 | 0.31 | 0.31 | 0.29 | 0.33 | 0.34 | 0.29 | 0.29 | 0.29 | 0.30 | 0.33 | 0.30 | 0.26 | | | |
| 17 | Umbelopsis dimorpha strain CBS 110039 | 0.30 | 0.28 | 0.33 | 0.31 | 0.31 | 0.29 | 0.33 | 0.34 | 0.29 | 0.29 | 0.29 | 0.30 | 0.33 | 0.30 | 0.26 | 0.00 | | |
| 18 | Bifiguratus adelaide NRRL 66559 | 0.37 | 0.36 | 0.47 | 0.45 | 0.43 | 0.37 | 0.42 | 0.43 | 0.36 | 0.34 | 0.34 | 0.40 | 0.43 | 0.40 | 0.49 | 0.45 | 0.45 | - |
