## Supplementary figures and images for "Pygmaeomycetaceae, a new root associated family in Mucoromycotina from the pygmy pine plains"

### SUPPLEMENTARY FIG. 1

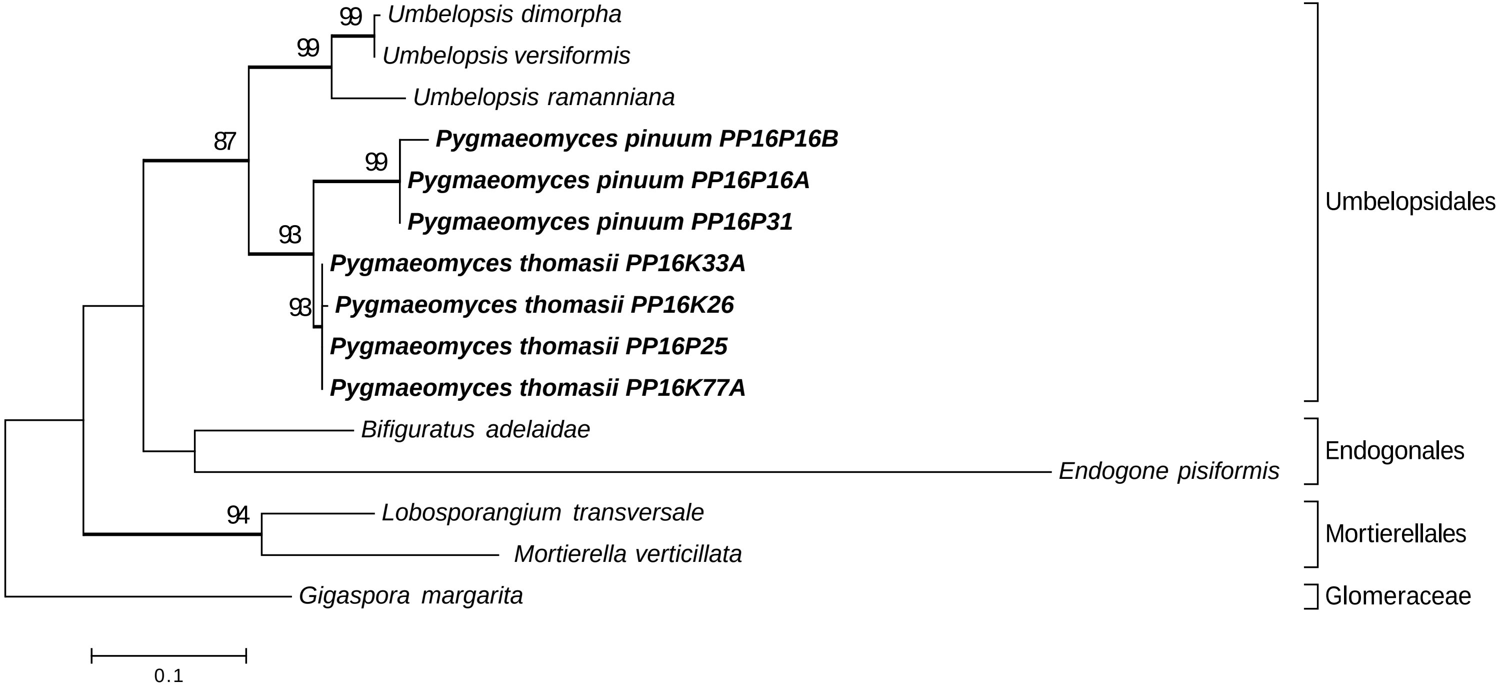

### SUPPLEMENTARY FIG. 2

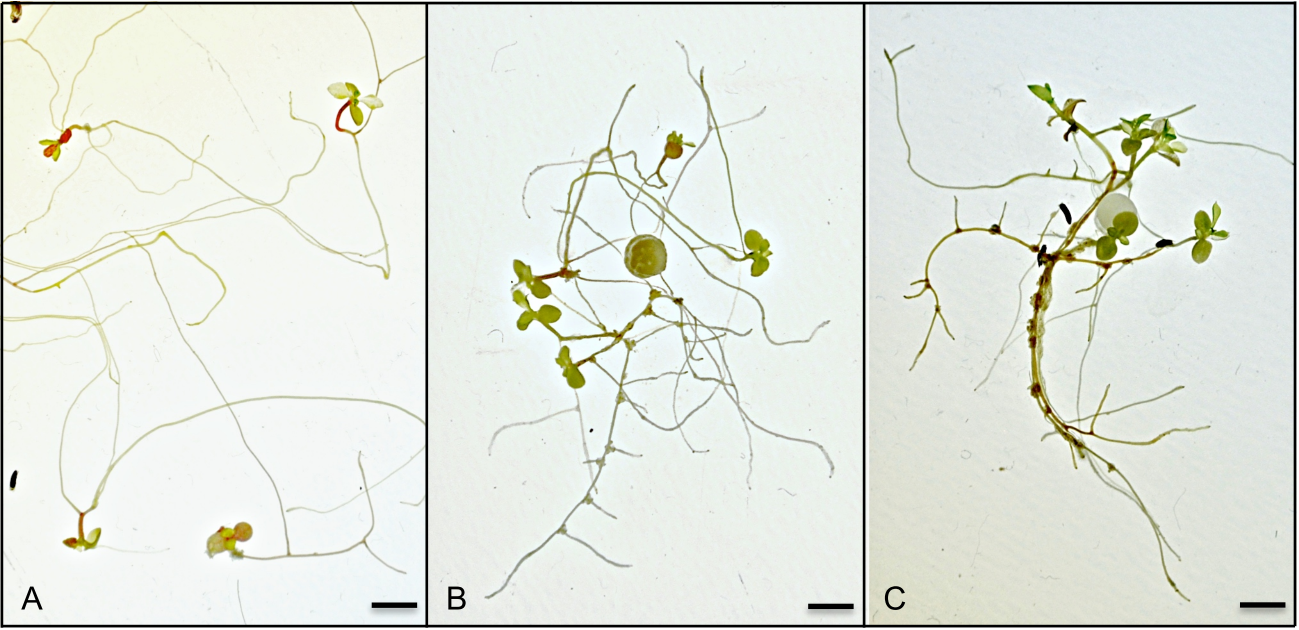
